# Copy Number Variation Analysis of 100 Twin Pairs Enriched for Neurodevelopmental Disorders

**DOI:** 10.1101/152611

**Authors:** Sofia Stamouli, Britt-Marie Anderlid, Charlotte Willfors, Bhooma Thiruvahindrapuram, John Wei, Steve Berggren, Ann Nordgren, Stephen W Scherer, Paul Lichtenstein, Kristiina Tammimies, Sven Bölte

**Affiliations:** Center of Neurodevelopmental Disorders (KIND), Department of Women’s and Children’s Health, Karolinska Institutet, Solna, Sweden; Center for Psychiatry Research, Stockholm County Council, Stockholm, Sweden; Department of Molecular Medicine and Surgery, Center of Molecular Medicine, Karolinska Institutet, Solna, Sweden; Department of Clinical Genetics, Karolinska University Hospital, Solna, Sweden; The Centre for Applied Genomics, Program in Genetics and Genome Biology, the Hospital for Sick Children, Toronto, Ontario, Canada; Department of Molecular Genetics and the McLaughlin Centre, University of Toronto, Toronto, Canada; Department of Medical Epidemiology and Biostatistics, Karolinska Institutet, Stockholm, Sweden

**Keywords:** Monozygotic twins, Neurodevelopmental disorders, Autism spectrum disorder, Copy number variant, Postzygotic variants, Discordant twin pairs

## Abstract

Hundreds of penetrant risk loci have been identified across different neurodevelopmental disorders (NDDs), and these often involve rare (<1% frequency) copy number variations (CNVs), which can involve one or more genes. Monozygotic (MZ) twin pairs are long thought to share 100% of their genomic information. However, genetic differences in the form of postzygotic somatic variants have been reported recently both in typically developing (TD) and in clinically discordant MZ pairs. Here, we sought to investigate the contribution of CNVs in 100 twin pairs enriched for NDD phenotypes with a particular focus on MZ pairs discordant for autism spectrum disorder (ASD) using the PsychChip array. In our collection, no postzygotic *de novo* CNVs were found in 55 MZ twin pairs, including the 13 pairs discordant for ASD. When analyzing the burden of rare CNVs among pairs concordant and discordant for ASD/NDD in comparison with typically developed (TD) pairs, no differences were found. However, we did detect a higher rate of CNVs overlapping genes involved in disorders of the nervous system in MZ pairs discordant and concordant for ASD in comparison with TD pairs (p=0.02). Our results are in concordance with earlier findings that postzygotic *de novo* CNV events are typically rare in genomic DNA derived from saliva or blood and, in the majority of MZ twins, do not explain the discordance of NDDs. Still, studies investigating postzygotic variation in MZ discordant twins using DNA from different tissues and single cells and higher resolution genomics are needed in the future.

## Introduction

Neurodevelopmental disorders (NDDs), such as autism spectrum disorder (ASD) and attention-deficit hyperactivity disorder (ADHD), are a group of early onset conditions affecting approximately 10-15% of the population.^1^ NDDs are characterized by alterations in developmental trajectories causing impairments in personal, social, academic, and occupational functioning.^2^ There is substantial co-occurrence of different NDDs suggesting a common etiology for the disorders.^3^ NDDs have a strong genetic component, and the familial risk is seen between different NDDs and psychiatric disorders.^4^ The etiological overlap has also been observed in molecular genetic studies.^5^ Many of the same risk genes and variants, identified through genome analyses of copy number variants (CNVs) and sequence level variants are found across NDDs. These include large deletions and duplications of 16p11.2 and 22q11.2, duplication of 7q11.23 and 15q11-13 as well as smaller CNVs affecting specific genes such as *NRXN1* and *SHANKs*. Additionally, new NDD risk genes have been implicated using sequencing-based studies mainly through identification of *de novo* mutations (DNMs). ^6^^-^^9^

In addition to germline DNMs, somatic DNMs contribute to human disorders^10^ including NDDs.^11,12^ It has been hypothesized that postzygotic DNMs could cause discordant phenotypes in monozygotic (MZ) twin pairs that otherwise are genetically identical. However, the majority of studies testing this hypothesis have reported no or limited evidence for the postzygotic *de novo* CNVs in MZ discordant twin pairs for psychiatric disorders^13^ and neurodevelopmental problems^14-16^. However, there are reports of postzygotic CNVs in MZ twin pairs without discordant phenotypes.^15,17^

Here, we analyze the contribution of CNVs in a Swedish twin sample from the Roots of Autism and ADHD Study in Sweden (RATSS)^18^. The sample is enriched for twin pairs discordant for NDDs including 13 MZ twin pairs discordant for ASD. Beyond rare case reports ^16^, there have not been studies focused on ASD-discordant MZ pairs and the contribution of postzygotic CNVs. We aimed to test if postzygotic *de novo* CNVs contribute to the discordancy for ASD and NDD phenotypes in MZ pairs in our sample. Additionally, we hypothesized that shared rare CNVs could increase the risk for NDDs even in discordant pairs. We investigated this hypothesis by analyzing the burden of rare CNVs and the rate of rare CNVs affecting genes earlier implicated in NDDs among MZ twin pairs concordant or discordant for NDD and ASD in comparison with typically developing (TD) twin pairs.

## Material and Methods

### Participants and clinical assessments

This study is approved by the National Swedish and the regional ethical review board at Karolinska Institutet and by the Swedish Data Inspection Board. Written informed consent was obtained from the twins (for twins older than 15 years) and/or their parents/legal guardians. The family identifiers have been randomized for this study and do not correspond to the identifiers assigned during the recruitment in the RATSS study.

All the participants included in this study are part of the RATSS.^18^ The recruitment and procedure for the sample collection have been described in details elsewhere.^18,19^ Overview of the study design and samples used for the current analyses is shown in eFigure 1. A total of 211 twins consisting of 104 twin pairs (a triplet was counted as one MZ female pair and one dizygotic (DZ) brother) were included, collected within RATSS between 2011 and 2014. In three families, two twin pairs were included. Additionally, 75 parents from 37 of these families were analyzed.

Within the RATSS, twins undergo detailed diagnostic and psychological assessments, including standardized diagnostic tools for NDDs; a psychosocial and anamnestic interview, the Autism Diagnostic Interview–Revised (ADI-R), the Autism Diagnostic Observation Schedule Second Edition (ADOS-2), the Kiddie Schedule for Affective Disorders and Schizophrenia (K-SADS) or the Diagnostic Interview for ADHD in Adults (DIVA). Additionally, the Wechsler Intelligence Scales for Children or Adults, Fourth Edition (WISC-IV) or the Leiter-revised scales in combination with the Peabody Picture Vocabulary Test, Third Edition and the parent-rated Adaptive Behavior Assessment Scale, 2nd Edition (ABAS-2), were used to evaluate adaptive, cognitive and verbal abilities and for diagnostic assessments. Also, a medical and family history for the twins was obtained by parental interviews and by collecting existing medical records.

The given NDD diagnoses are based on the DMS-5 criteria^2^ by incorporating the different assessments followed by a consensus between three experienced clinicians. Using the Child and Adolescent Twin Study in Sweden (CATSS) cohort, which is also a recruitment cohort for RATSS, it has been demonstrated that in the majority of ASD-discordant MZ pairs, the co-twin has another NDD diagnosis.^20^ Therefore, in this study, we used two strategies to categorize the twins based on their NDD diagnoses. First, we only considered if any of the twins had ASD diagnosis and categorized the pair as discordant or concordant for ASD. Thereafter, we considered all NDD diagnoses and did an additional categorization as NDD discordant (affected twin has any NDD diagnosis and co-twin TD) and NDD concordant (both twins have at least one NDD diagnosis). If no NDD diagnoses were present in any of the twins, the pair was categorized as TD.

### DNA samples and genotyping

Saliva samples were collected from the twins and their parents using the Oragene·DNA OG-500 tubes (DNA Genotek, Inc., Ottawa, Ontario, Canada) during the RATSS study visit or at home. The extraction of genomic DNA from the saliva samples was done automatically using the Autopure LS system and Puregene DNA purification kit (Gentra Systems, Minneapolis, MN, USA) or Chemagen DNA saliva kit (PerkinElmer, Waltham MA, USA). A total of 200 ng of DNA was used for the genotyping using the Infinium PsychArray-24 v1.1 (Illumina Inc., San Diego, California, USA) according to the manufacturer’s protocol. Samples with SNP call rate lower than 98% were excluded from further investigations. The zygosity of the twin pairs was confirmed using the genotype data by estimating the identity by descent using the PLINK/1.07^21^ after quality control and removal of SNPs with minor allele frequency <0.05 within the samples.

### CNV calling

We derived the CNVs in our data by using four different algorithms: PennCNV^22^, QuantiSNP^23^, iPattern^24^ and iPsychCNV (pre-release beta v1.0). For the downstream analyses, we used only stringent CNVs, which were called by iPattern and at least one additional algorithm or by PennCNV together with iPattern or/and iPsychCNV with 50% reciprocal overlap between the algorithms. We further excluded variants that were <10 kb in size, cover by <10 consecutive probes and had more than 50% overlap with centromeres. After obtaining the final set of stringent CNV calls, we removed individuals with excessive CNV calls defined by >2 standard deviations from the mean CNV count in the whole sample. Population-based control data sets described by Lionel *et al*.^25^ were used to restrict the analyses to rare CNVs found in frequency <0.01% in controls. Additionally, we computed the frequency of CNV events using PLINK/1.07^21^ based on two reference sets; the total cohort including both parents and the twins, and the parents only. We removed variants present > 3 in the parents.

### Pre-twinning and post-twinning de novo CNV detection

The pre-twinning *de novo* CNVs were analyzed in 18 families in which DNA was available for both parents. The rare stringent CNVs shared by the twin pair or present in one twin in the twin pairs and not found in the parents or the whole population stringent and non-stringent CNV calls were considered as putative pre-twinning *de novo* variants. Similarly, the postzygotic *de novo* CNV events were analyzed using the stringent CNV set from the 55 complete MZ twin pairs. Rare CNVs found only in one twin and not detected in any of the calls of the population dataset were considered as post-twinning *de novo* variants. The putative pre- and post-twinning *de novo* CNV calls were visually inspected by plotting the Log R Ratio (LRR) and B allele frequency values of the twins and the parents.

### Burden of rare CNVs and CNVs overlapping specific set of genes

Next, we computed rare CNV characteristics for the MZ and DZ twins groups based on the concordance/discordance for ASD and NDD. We included all MZ twin pairs for which at least one twin was successfully genotyped and each twin in DZ pairs. Additionally, we identified CNVs that overlapped known genomic disorders, earlier implicated in ASD/intellectual disability genes^26^, genes curated in the SFARI gene database (accession date September 2016)^27^ and genes associated with abnormality of the nervous system in the human phenotype ontology (HP:0000707) hereafter termed as HPO-NS^28^. Thereafter the CNVs were also evaluated based on the criteria provided by the American College of Medical Genetics and Genomics (ACMG)^29^.

We tested the differences in the rare CNV characteristics among the diagnostic groups in MZ pairs as well as between twins in discordant DZ pairs. For these groups, the following CNV characteristics were compared: the average total rare CNV length, the average rare CNV length, the average number of genes with coding sequence (CDS) affected by rare CNVs, the average rare CNV length overlapping CDS, the rate of rare CNVs and the rate of rare CNVs overlapping CDS. The statistical differences in the length and number of genes were tested using Kruskal-Wallis test within the MZ twin pairs. Additionally, three-group χ^2^ or Fisher’s exact test (FET) were applied to assess if the rate of rare CNVs and the rate of rare CNVs overlapping CDS genes per twin pair were significantly different in the MZ twin categories.

In DZ pairs, the intra-pair differences in the rare CNVs characteristics among twins discordant for ASD/NDD were tested using Wilcoxon signed rank test. McNemar’s test was used to compare the rate of rare CNVs and the rate of rare CNVs affecting CDS among the DZ discordant pairs. The TD pairs and the pairs concordant for ASD/NDD were not used in the statistical tests due to their small sample size. All statistical analyses were performed with R (3.2.3).

### Experimental validation of selected CNVs

Putative pre- and post-zygotic *de novo* and NDD associated CNVs that passed our filtering criteria and visual inspection were subjected to experimental validation using quantitative PCR (qPCR). For this, Taqman^®^ probes (Thermo Fisher Scientific, Waltham MA, USA) within the genomic region of interest for the validation experiment were chosen and tested together with a reference control probe targeting *TERT* gene (Thermo Fisher Scientific). Copy number state of each region was tested in the twin pair, parental DNA if available and at least one unrelated control DNA sample. Quadruplicates of each sample were analyzed on StepOneTM (Thermo Fisher Scientific) system according to manufacturer’s recommendations. The copy number of the target in each sample was analyzed using the CopyCaller™ software (Thermo Fisher Scientific).

## Results

### Sample characteristics

Out of the 284 individuals genotyped, a total 231 were preceded for further analysis after the QC of genotyping and CNV calling (eFigure 1). The majority of the samples that failed QC were parental saliva samples collected at home. The final set comprised of 69 MZ (including one female MZ pair from the triplet) and 31 DZ twin pairs (the male sibling from the triplet was excluded from the analysis), corresponding to 97 families. For 55 MZ and 19 DZ pairs, both twins were successfully genotyped. In 18 families both parents were genotyped. The twins were classified into twin pairs discordant or concordant for ASD and NDD based on the DSM-5 diagnoses assigned within the study. A total of 26 pairs were discordant and 11 pairs concordant for ASD (Table 1). In the twin pairs discordant for ASD, 69.2% (18/26) of the affected twins and 19.2% (5/26) of the co-twins had at least one additional NDD diagnosis. For the NDD category, 34 pairs were classified as discordant and 35 pairs as concordant. Twenty-seven twin pairs were classified as typically developed. Mean age, IQ, and the number of other DSM-5 NDD diagnoses for the twin pairs in each category are shown in Table 1.

**Table 1.** Phenotypic characteristics of the monozygotic and dizygotic twins in the study

| *MZ twin pairs* | *Typically developed* | *Concordant ASD* | *Discordant ASD* |  | *Concordant NDD* | *Discordant NDD* |  |
| --- | --- | --- | --- | --- | --- | --- | --- |
| *Number of pairs* | 25 | 10 | 13 |  | 22 | 19 |  |
| *Female pairs (%)* | 12 (48%) | 5 (50%) | 5 (38.5%) |  | 7 (31.8%) | 9 (47.4%) |  |
| *Male pairs (%)* | 13 (52%) | 5 (50%) | 8(61.5%) |  | 15 (68.2%) | 10 (52.6%) |  |
| *Age mean (range)* | 15.6 (10-22) | 15.5 (9-28) | 14 (9-20) |  | 15 (9-28) | 13 (9-20) |  |
|  |  |  | **Affected** | **Co-twin** |  | **Affected** | **Co-twin** |
| *IQ mean (range)* | 101.8<br>(81-126) | 88.1<br>(58-142) | 88.3<br>(40-138) | 96.8<br>(65-121) | 86.7<br>(58-142) | 93.8<br>(40-138) | 98.8<br>(77-121) |
| *Cases with ASD (%)* | - | 20 (100%) | 13 (100%) | - | 20 (45.5%) | 10 (52.6%) | - |
| *Cases with ADHD (%)* | - | 11 (55%) | 2 (15.4%) | 1 (7.7%) | 28 (63.6%) | 9 (47.4%) | - |
| *Cases with ID (%)* | - | 3 (15%) | 3 (23.1%) | 1 (7.7%) | 11 (25%) | 3 (15.8%) | - |
| *Cases with other NDD diagnoses \** | - | 4 (20%) | 1 (7.7%) | - | 15 (34.1%) | 6 (31.6%) | - |
| *DZ twin pairs* | *Typically developed* | *Concordant ASD* | *Discordant ASD* |  | *Concordant NDD* | *Discordant NDD* |  |
| *Number of pairs* | 2 | 1 | 13 |  | 13 | 15 |  |
| *Female pairs (%)* | - | - | 2 (15.4%) |  | 5 (38.5%) | 5 (33.3%) |  |
| *Male pairs (%)* | 1 (50%) | 1 (100%) | 8 (61.5%) |  | 7 (53.8 %) | 7 (44.7%) |  |
| *Opposite sex pairs (%)* | 1 (50%) | - | 3 (23.1%) |  | 1 (7.7%) | 3 (20%) |  |
| *Age mean (range)* | 13.5 (11-16) | 14.0 (14-14) | 14.1 (10-20) |  | 14.2 (8-19) | 14.6 (8-25) |  |
|  |  |  | **Affected** | **Co-twin** |  | **Affected** | **Co-twin** |
| *Mean IQ (range)* | 125.0<br>(115-138) | 97.0<br>(83-111) | 83.5<br>(42-120) | 98.2<br>(65-124) | 95.3<br>(65-130) | 88.1<br>(42-120) | 101.9<br>(77-124) |
| *Cases with ASD (%)* | - | 2 (100%) | 13 (100%) | - | 2 (7.7%) | 10 (66.7%) | - |
| *Cases with ADHD (%)* | - | 1 (50%) | 8 (61.5%) | 2 (15.3%) | 20 (77%) | 10 (66.7%) | - |
| *Cases with ID (%)* | - | - | 3 (23.1 %) | 1 (7.7%) | 2 (7.7%) | 2 (13.3%) | - |
| *Cases with other NDD diagnoses\** | - | - | 1 (7.7%) | - | 3 (11.5%) | 4 (26.7%) | - |

\*Additional NDD diagnosis include specific learning disorder, tic disorder, and speech sound disorder

### Pre-twinning de novo CNVs

We analyzed pre-twinning *de novo* CNVs in 18 families in which both parents were successfully genotyped. Two putative *de novo* CNVs were identified in two families (eTable 1). However, none of these were validated to be *de novo* by qPCR. One CNV was false positive, and a ~85 kb duplication affecting *CPA6* shared by MZ twin pair discordant for ADHD was shown to be maternally inherited.

### Post-twinning de novo CNVs

We aimed to identify putative postzygotic *de novo* CNV events in the 55 complete MZ twin pairs of which 11 pairs were classified as discordant for ASD or 18 pairs as discordant for NDDs. First, we calculated the concordance rate of each CNV detection algorithm when using the <10 kb size and <10 consecutive probe filter within the twin pairs (eTable 2). The average concordance rates ranged between 13.6% for iPsychCNV to 65.3% for iPattern. In our final stringent CNV calls, the concordance rate was 64.2% with 16 of 55 twin pairs (29.1%) sharing all their CNVs. A total of eight non-shared rare CNV calls were found in four MZ twin pairs in our analysis after our filtering procedure. After visual inspection, five proceeded to qPCR validation. None of the five CNVs were validated to be non-shared; two were found in both twins, and three were a false positive (eTable1).

### Burden of rare CNVs

Next, we tested if there were significant differences in the rate of rare CNVs and the rate of rare CNVs overlapping coding sequence between the TD twin pairs and discordant or concordant for ASD/NDD. Additionally, we examined the difference in the average total length, the average length, the average number of genes within the region of rare CNVs among the twin pair groups (eTable 3, eTable 4). We did not find any statistically significant differences among the MZ twin pairs in the different phenotypic categories.

Similarly, we tested if any differences for the CNV characteristics were found between the affected and the co-twin in DZ twins discordant for ASD and/or NDD (eTable 4). Within the ASD-discordant DZ pairs, we show that the affected twin has increased average total length and the average length of rare CNVs (p=0.03 and p=0.04, respectively).

### Rare CNVs affecting genes previously implicated in NDDs or nervous system disorders

In total, 28 CNVs in 22 twin pairs were found to overlap the predefined gene sets, including ASD/ID genes^26^ in combination with SFARI genes^27^ and/or HPO-NS genes. We calculated the proportion of twin pairs or twins in discordant DZ pairs having at least one CNV in these gene categories within each twin category. We found an increased rate of MZ twin pairs discordant and concordant for ASD having CNVs affecting both NDD genes (Figure 1A) and HPO-NS (Figure 1B) in comparison with TD MZ pairs. However, only the HPO-NS comparison reached statistical significance (3-group comparison 2- sided FET p=0.02). As expected the largest difference was found between the frequency in TD pairs compared with concordant ASD pairs (OR=14.4, 95% CI 1.2-814.6). For MZ twins categorized based on the NDD diagnosis, we did not observe any significant differences for the NDD or HPO-NS (3-group comparison, 2-sided FET p=0.92, and p=0.11, respectively). However, there was a minor increase in the proportion of twin pairs with CNVs for both NDD concordant and discordant pairs in comparison with the TD pairs for the HPO-NS genes (Figure 1D). For DZ twin pairs, no differences were found between ASD/NDD affected twins and their co-twins (eTable 5).

**Figure 1.**
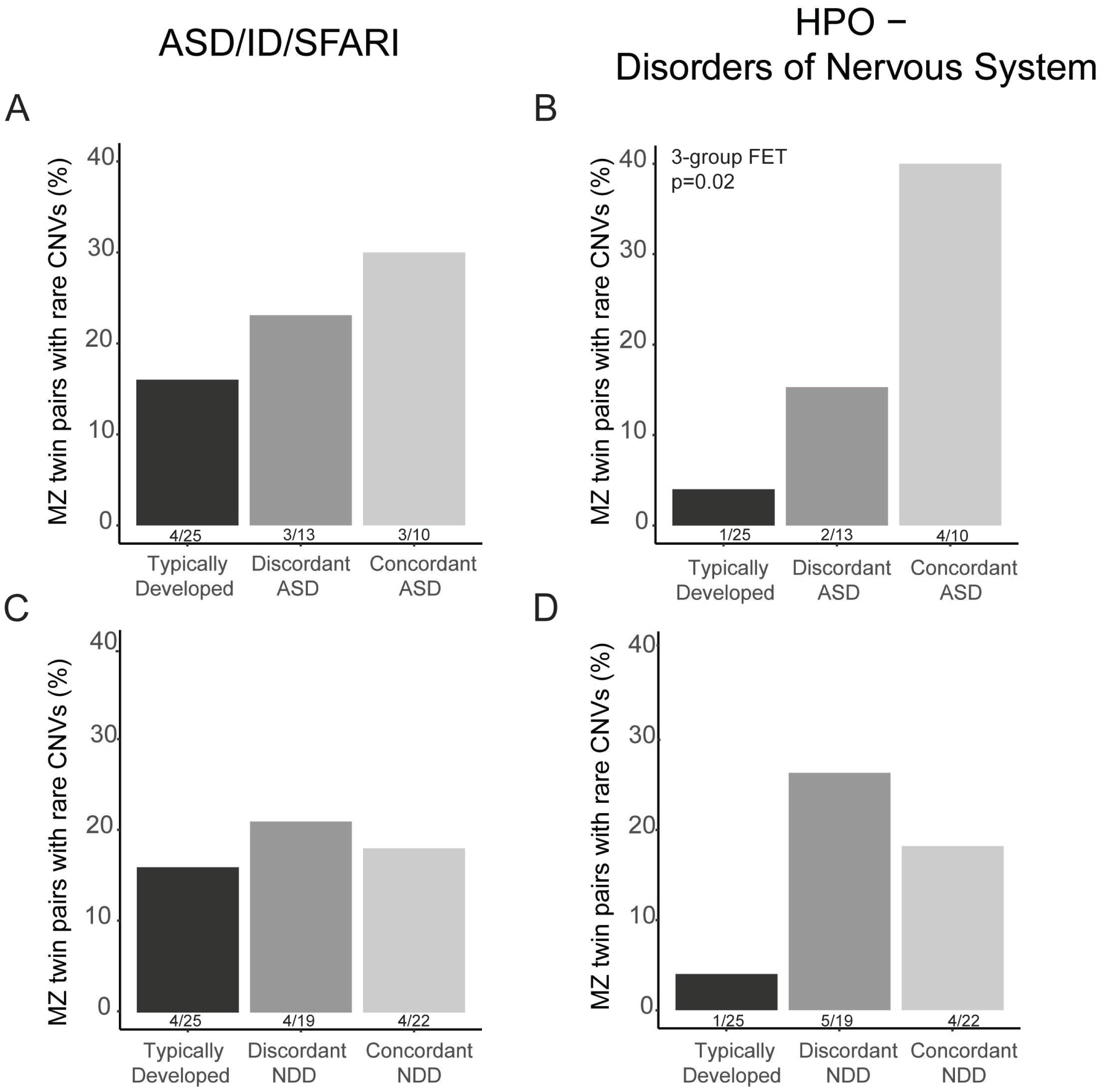
The rate of rare CNVs affecting genes earlier implicated in neurodevelopmental disorders (ASD/ID/SFARI) or abnormality of the nervous system in the Human Phenotype Ontology database (HPO-NS) in monozygotic twin pairs. The percentage of MZ twins in typically developing, ASD-discordant and –concordant pairs with CNVs overlapping genes in ASD/ID/SFARI gene lists (A) and in HPO-NS (B). The percentage of typically developing, NDD-discordant and -concordant pairs with CNVs overlapping genes in ASD/ID/SFARI gene lists (C) and in HPO-NS (D). Statistical significance was tested between the three twin pair groups using Fisher Exact Test, p-value for significant comparison is reported in the figure.

Next, we investigated closer the rare CNVs overlapping the ASD/ID and HPO-NS gene lists to categorize them based on the guidelines from the ACMG ^29^ and evidence from the earlier literature. Two CNVs in two twin pairs among the 10 MZ pairs concordant for ASD (20.0%) and one additional CNV among the other MZ pairs concordant for NDD (13.6%, 3 out of 22 pairs), were classified as clinically significant. A male MZ twin pair concordant for ASD and ADHD had a paternally inherited ~9.0 Mb duplication at 12q11-12q13.11 affecting 33 genes. Proximal deletions of 12q12 had earlier been implicated in patients with NDDs and dysmorphic features^30^. Another MZ male pair concordant for ASD and ID and discordant for tic disorder had a ~44.3 kb deletion on chromosome 1q44 affecting *COX20* and *HNRNPU*, as well as *HNRNPU-AS1* long-coding RNA (Figure 2). We were not able to conclude if the deletion was of de novo origin as paternal DNA sample was not available. Recently, two reports described case series with mutations affecting *HNRNPU* demonstrating its role in severe intellectual disability and early-onset seizures. ^31^ Also, recent exome studies for developmental delay^32^ and ASD^33^ have identified eight loss-of-function and two missense *de novo* mutations affecting *HNRNPU* (Figure 2). A maternally inherited 18.8 kb deletion on chr8p23.2 disrupting *CSMD1* was detected in MZ pair concordant for ADHD and discordant for tic disorder. *CSMD1* has been implicated in schizophrenia and cognitive ability through association studies. ^34,35^ Recently, both inherited and *de novo* variants in *CSMD1* have been found in ASD patients.^33,36^

**Figure 2.**
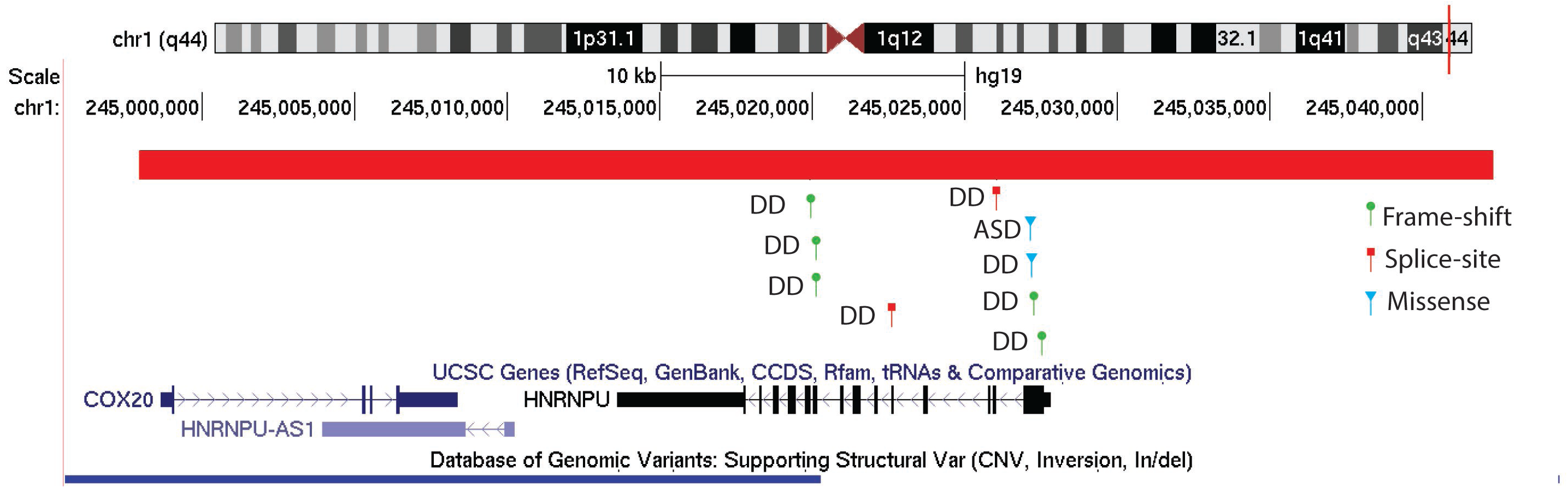
Schematic of the *HNRNPU* genomic region on Chr1q44. The deletion identified in a male monozygotic twin pair concordant for intellectual disability and autism spectrum disorder is highlighted in red. A collection of disrupting *de novo* mutations within the *HNRNPU* gene found in developmental delay (DD)^29^ and autism spectrum disorder (ASD)^30^ from the literature are highlighted below the deletion. The blue vertical box represents a duplication within the region in the Database of Genomic Variants.

Among the discordant MZ pairs, we detected a maternally inherited 0.95 Mb gain on chromosome 6q13-14.1 possible contributing to the phenotypes seen in the male twin pair. Both large deletions and duplications at chromosome 6q encompassing this region have been implicated in NDD associated phenotypes.^37,38^ The affected twin has ASD with full-Scale IQ of 70, dyslexia, and earlier indicated mild macrocephaly (current head circumference 54.5cm, 84^th^ percentile). The co-twin has mild macrocephaly (56 cm, 98th percentile) and a reported mild language delay described as language impairments specific for ASD, i.e. reversal use of pronouns and regularly use of idiosyncratic language, as well as mild understanding and articulation problems. Lack of social reciprocity, pretend, and fantasy play is reported for both twins. However, the impairments in the co-twin are overall reported as previous during development and at a sub-threshold level, and not observed at the time of assessment, in comparison to the affected twin. The co-twin also has a slightly higher IQ of 77.

In the DZ pairs, we identified a CNV affecting *SYNE1* that is likely pathogenic in a male twin diagnosed with ASD and ID from an opposite-sex DZ pair. We verified the 39 kb gain on chr6q25.2 using qPCR and detected seven copies of the region in the affected male twin (eFigure 2). His TD female sibling and mother both had normal CN for the region; no DNA sample was available for testing from the father. *SYNE1*, which encodes for the Nesprin-1 protein, has been implicated in ASD through rare mutations^33,39,40^ but is also involved in spinocerebellar ataxia^41^ and Emery-Dreifuss muscular dystrophy^42^.

## Discussion

Our aim was to investigate the presence of postzygotic *de novo* CNVs that could explain the discordance of a variety of NDD phenotypes, especially ASD, in MZ pairs, and to test if the burden of rare CNVs was different between the twin pairs in our sample of 100 twin pairs based on their concordance of NDDs. We did not identify any validated postzygotically derived unique CNVs in the 55 MZ pairs analyzed, thus indicating that these are not major contributors for the discordancy of NDDs in our sample nor a common phenomenon among MZ twin pairs. Our results are in accordance with the earlier studies showing that postzygotic *de novo* CNVs are rare when analyzing genomic DNA from blood or saliva samples.^13-17^ As we only restricted our analysis to saliva DNA and CNV level variants, we cannot exclude the possibility that other types or tissue-specific postzygotic somatic variations contribute to the discordancy. In non-twin cohorts, somatic sequence level mutations have been shown to contribute to both ASD and ID diagnoses. ^12,43,44^ Additionally, somatic CNVs and sequence levelvariation can be found in human neurons from healthy individuals using single cell genomics.^45,46^ Therefore, stochastic genetic variations at a neuronal level could contribute to disrupted brain development and underlie phenotype differences connected to brain disorders in MZ twins. As recent literature has implicated somatic DNMs have a role in NDDs, a better understanding of their contribution and origin in NDDs is needed as it can inform better estimations of the recurrence rates of the disorders and thus genetic counseling.

We also sought to understand the contribution of shared rare CNVs in the twin pairs with a specific focus on the ASD and NDD-discordant twins. As expected, we identified a higher rate of rare CNVs, especially in genes earlier implicated in NDDs and development of the nervous system in ASD-concordant pairs (Figure 1). The rate of putatively clinical significant CNVs in our twins concordant for NDD (13.6 %) was similar to the reported rates in ASD cohorts ranging from ~10% to 28%.^9,47^ Interestingly, we detected an increased rate of the CNVs affecting genes involved in alterations of the nervous system in the ASD/NDD-discordant MZ pairs (Figure 1B, D) demonstrating that CNVs with incomplete penetrance could contribute to the NDD phenotypes seen in the twins. Many of the most common CNVs found in NDD cases have variable expressivity as well as incomplete penetrance.^48^ More severe phenotypes connected to the recurrent genomic disorders and rare CNVs are most likely due to additional genetic variants, epigenetic modifications, environmental exposures and their combined effects. For instance, additional rare CNVs were more common in carriers of the recurrent 16p11.2 deletions with more severe NDD phenotypes.^49^ The benefit of MZ discordant twin pairs design is the possibility to pinpoint these additional factors such as non-shared environmental factors that contribute to the etiology of NDDs alone or as modifying the effect of rare CNVs. Indeed, we have identified non-shared medical and environmental factors that are contributing to the phenotypes using this same sample of twins.^19,50^

Multiple studies have highlighted the need for a cross-disorder approach in genetic studies as earlier research have shown the clinical and genetic overlap between different NDDs.^3,51^ We used two different approaches, either clinical ASD diagnosis only or the umbrella category of NDD, to define the concordance and discordance of the twins for our analysis of shared CNVs. Using this approach, we demonstrate a rising degree of the rate of CNVs affecting specific genes from TD to ASD-discordant and -concordant pairs. When using the NDD concept, we did not find any differences among the NDD discordant and concordant pairs despite an elevated rate seen in comparison with the TD pairs. The lack of difference could be due to the milder phenotypes such as learning disabilities included in the concordant NDD group that will lower the proportion of more severely affected twin pairs in this group. It has been implicated that CNV burden is not increased in milder NDDs.^52^

In conclusion, we add to the growing literature that postzygotic *de novo* CNV events in MZ twin pairs are rare and do not explain the discordance of NDD phenotypes in the majority of the twin pairs. We suggest that shared CNVs overlapping genes essential for brain development increase the risk for NDDs even in discordant MZ pairs. Further studies using larger twin cohorts, additional sources of DNA, and better variant detection resolution such as whole genome sequencing^8^ are needed to confirm our results and overcome the limitations of this study. Additionally, studies should aim at investigating the contribution of other risk or resilience factors contributing to the phenotypic outcomes in genetic risk backgrounds.

### Authors’ contributions

S.S., S.Bö., and K.T, conceived and designed the study. B-M.A., C.W., S.Be., A.N. recruited, diagnosed and examined the twins. S.S., B.T., J.W., and K.T. processed the microarray data. S.S. and K.T. analyzed the data and drafted the manuscript. All authors contributed to acquisition of data, interpretation of results and critical discussion and approved the final version of the manuscript.

## Acknowledgements

We thank all the twins and their families for participating in this study. We also thank Kerstin Andersson, Christina Coco, Martin Hammar, Johanna Ingvarsson, Anna Lange Nilsson, Gunnel Ahréns, Lynnea Myers, Lina Poltrago and Anna Råde, for their valuable contribution to the work presented in this study. Genotyping was performed by the SNP&SEQ Technology Platform in Uppsala (www.genotyping.se). The facility is part of the National Genomics Infrastructure (NGI) Sweden and Science for Life Laboratory. The SNP&SEQ Platform is also supported by the Swedish Research Council and the Knut and Alice Wallenberg Foundation. The computation resources were provided by SNIC through Uppsala Multidisciplinary Center for Advanced Computational Science (UPPMAX) (b2015257).

Support was provided by the Innovative Medicines Initiatives Joint Undertaking (grant agreement number 115300), which comprises financial contribution from the European Union’s Seventh Framework Programme (FP7/2007 – 2013) and in-kind contributions from companies belonging to the European Federation of Pharmaceutical Industries and Associations; the Swedish Research Council (523-2009-7054; 521-2013- 2531; 350-2012-286); the Swedish Research Council, in partnership with the Swedish Research Council for Health, Working Life and Welfare, Formas and VINNOVA (cross-disciplinary research program concerning children’s and young people’s mental health, 259-2012-24), Stockholm County Council (20100096, 20110602, 20120067, 20140134), Stiftelsen Frimurare Barnhuset, Sunnerdahls-Handikappfond, Hjärnfonden. S.W.S. is supported by the GlaxoSmithKline-Canadian Institutes of Health (CIHR) Endowed Chair in Genome Sciences at The Hospital for Sick Children and University of Toronto. K.T is financially supported by the Swedish Foundation for Strategic Research, the Harald and Greta Jeanssons Foundations, Åke Wiberg Foundation, The Swedish Foundation for International Cooperation in Research and Higher Education, StratNeuro, The Board of Research at Karolinska Institutet and Foundations and Funds at Karolinska Institutet.

## Conflict of interest

Sven Bölte declares no direct conflict of interest related to this article. He discloses that he has in the last five years acted as an author, consultant or lecturer for Shire, Medice, Roche, Eli Lilly, Prima Psychiatry, GLGroup, System Analytic, Kompetento, Expo Medica, and Prophase. He receives royalties for textbooks and diagnostic tools from Huber/Hogrefe, Kohlhammer, and UTB. Stephen Scherer is on the Scientific Advisory Boards of Population Diagnostics, Deep Genomics, DNAStack, and has intellectual property licensed relevant to CNVs through the Hospital for Sick Children to Lineagen. The other authors declare that they have no competing interests.

Supplementary information is available at the European Journal of Human Genetics website.

## Availability of raw data

The raw datasets generated and analyzed during the current study are not publicly available because they contain private patient health information, but are available from the corresponding author on reasonable request and subject to necessary clearances.

## Supplementary information (SI)

SI_file1 (docx) includes the following items:

**eFigure 1:** Overview of the study including the numbers of monozygotic (MZ) and dizygotic (DZ) pairs included in each step of the analysis.

**eFigure 2:** Visualization and validation of 39 kb gain on the chr6q25.2 present in a male twin (twin 2) with autism spectrum disorder and intellectual disability.

**eTable 1:** Predicted shared and postzygotic *de novo* copy number variants in the sample.

**eTable 2:** Copy number variant concordance of the different variant calling algorithms and the stringent CNVs used for the study.

**eTable 5:** The number of rare copy number variants overlapping genes implicated in the abnormality of the nervous system in the Human Phenotype Ontology (HPO-NS) and genes earlier implicated in neurodevelopmental disorders in dizygotic (DZ) discordant twin pairs (see Methods).

SI_file2 (Excel file) contains

**eTable 3:** All rare stringent copy number variants in the study samples after quality control.

**eTable 4:** Characteristics of rare copy number variants in monozygotic (MZ) and dizygotic (DZ) twin pairs

